# Fluorescent Detection of *O*-GlcNAc via Tandem Glycan Labeling

**DOI:** 10.1101/2020.06.27.175075

**Authors:** Zhengliang L. Wu, Ang Luo, Alex Grill, Taotao Lao, Yonglong Zou, Yue Chen

## Abstract

*O*-GlcNAcylation is a reversible serine/threonine glycosylation on cytosolic and nuclear proteins that are involved in various regulatory pathways. However, the detection and quantification of *O*-GlcNAcylation substrates have been challenging. Here we report a highly efficient method for the identification of *O*-GlcNAc modification via tandem glycan labeling, in which *O*-GlcNAc is first galactosylated and then sialylated with a fluorophore-conjugated sialic acid residue, therefore enabling highly sensitive fluorescent detection. The method is validated on various proteins that are known to be modified by *O*-GlcNAcylation including CK2, NOD2, SREBP1c, AKT1, PKM and PFKFB3, and on the nuclear extract of HEK293 cells. Using this method, we then report the evidence that hypoxia-inducible factor HIF1α is a target for *O*-GlcNAcylation, suggesting a potential direct connection between the metabolic *O*-GlcNAc pathway and the hypoxia pathway.

---

*O-*GlcNAc refers to a single β-*N-*acetylglucosamine residue attached to serine/threonine residues on nuclear and cytosolic proteins ^1^. *O-*GlcNAc is introduced by *O-*GlcNAc transferase (OGT) ^2^ from the donor substrate UDP-GlcNAc that is synthesized via the hexosamine biosynthetic pathway^3^. *O*-GlcNAc is removed by *O-*β-N-acetylglucosaminidase (OGA)^4^. *O-*GlcNAcylated proteins are primarily found in nuclei and secondary in cytoplasm^5^. Many intracellular proteins that are involved in transcriptional regulation are known to be targets of *O-*GlcNAcylation^6, 7^. For example, NOD2^8^ and SREBP-1^9^ are regulated by *O*-GlcNAcylation. NOD2 is a cytoplasmic innate immune receptor that upon activation can activate nuclear factor kappa B (NF-κB)^10^. SREBP-1 is a transcription factor that binds to sterol regulatory elements and regulates lipid metabolism^11^. Some intracellular enzymes that play key regulatory roles are also targets of *O*-GlcNAcylation. Such known enzymes include casein kinase II (CK2)^12^ and AKT1^13^ that are involved in regulations of cell metabolism, proliferation, survival, growth, and angiogenesis. One speculative *O*-GlcNAcylation target is a hypoxia-inducible factor (HIF), a master regulator of cellular and developmental response to hypoxia^14^. HIF consists of a subunit of HIF1α or HIF2α and a subunit of HIF1β. HIF pathway is known to be regulated by *O*-GlcNAcylation^15^, but it is not clear whether HIF itself is modulated by *O*-GlcNAcylation.

To study the biological functions of *O*-GlcNAc, methods for its specific and sensitive detection are critical. In addition to the classical antibody-based methods of detection^16^, many other types of methods have been explored for *O-*GlcNAc detection^17^, which usually involve an extension or metabolic labeling of *O*-GlcNAc with radioisotope^18^ 19 or chemically modified sugar residues ^20-24^. These methods are either complicated to perform or lacks specificities or sensitivity. Recently, we reported methods of direct fluorescent glycan labeling (DFGL) for detecting the substrate glycans of sialyltransferases ^25^ and fucosyltransferases ^26^. However, these methods cannot be directly applied to detect *O*-GlcNAc, as *O*-GlcNAc is not a direct substrate of any of these enzymes. Nevertheless, *O*-GlcNAc is known as a substrate for the β1,4 galactosyltransferase B4GalT1 with the product of *O*-linked lactosamine Galβ-1,4-GlcNAc-O-S/T ^18^. Since lactosamine can be labeled with fluorophore-conjugated sialic acids by the α-2,6-sialyltransferase ST6Gal1 ^25^, it is surmised that *O*-GlcNAc may be indirectly detected via a tandem labeling approach with both B4GalT1 and ST6Gal1 (Fig. 1).

**Figure 1.**
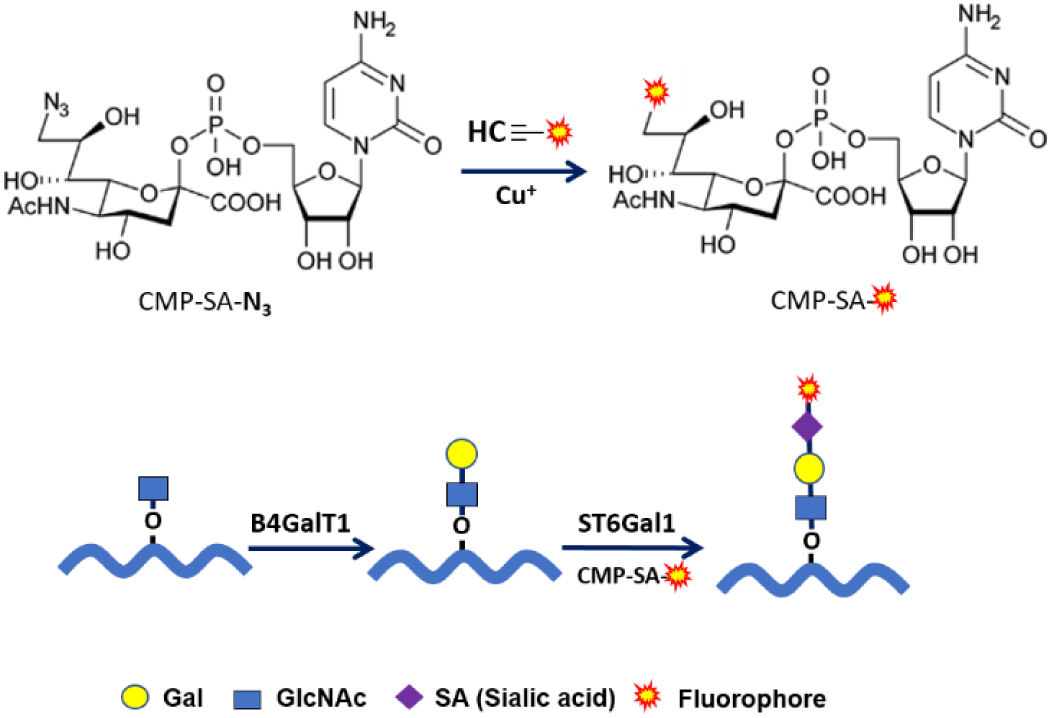
Fluorescent Detection of *O*-GlcNAc via Tandem Glycan Labeling. Fluorophore-conjugated activated sialic acids were prepared by copper (I)-catalyzed azide-alkyne cycloaddition (Cu-AAC)^27^ of azido-sialic acid and alkyne-fluorophores. O-GlcNAc is first galactosylated by B4GalT1 to become an *O*-linked lactosamine structure and then labeled with fluorophore-conjugated sialic acids by sialyltransferases, such as ST6Gal1.

To test the hypothesis proposed in Fig.1, the recombinant human CK2 (rhCK2) (Supplemental Fig. 1) was first successively treated by OGT, B4GalT1, and ST6Gal1 in the presence of their natural donor substrates. When the samples were separated on SDS-PAGE, successive mobility shift was observed (Fig. 2A), suggesting that the modifications did take place step by step. When the reactions were repeated and analyzed by mass spectrometry, the stepwise additions of *O*-GlcNAc, Gal, and sialic acid residues to rhCK2 were confirmed (Supplemental Fig. 2). The results of gel analysis and mass spectrometry analysis consistently suggest that the successive modifications on *O*-GlcNAc by B4GalT1 and ST6Gal1 are highly efficient and *O*-GlcNAcylation by OGT is rate-limiting. The hypothesis proposed in Fig.1 was further proved on rhCK2 by replacing the natural sialic acid with Alexa Fluor 488, Cy5, and Alexa Fluor 555-conjugated sialic acids. Fig. 2B shows the successive mobility shift caused by OGT, B4GalT1, and ST6Gal1 on rhCK2 and the incorporation of these fluorophore-conjugated sialic acids.

**Figure 2.**
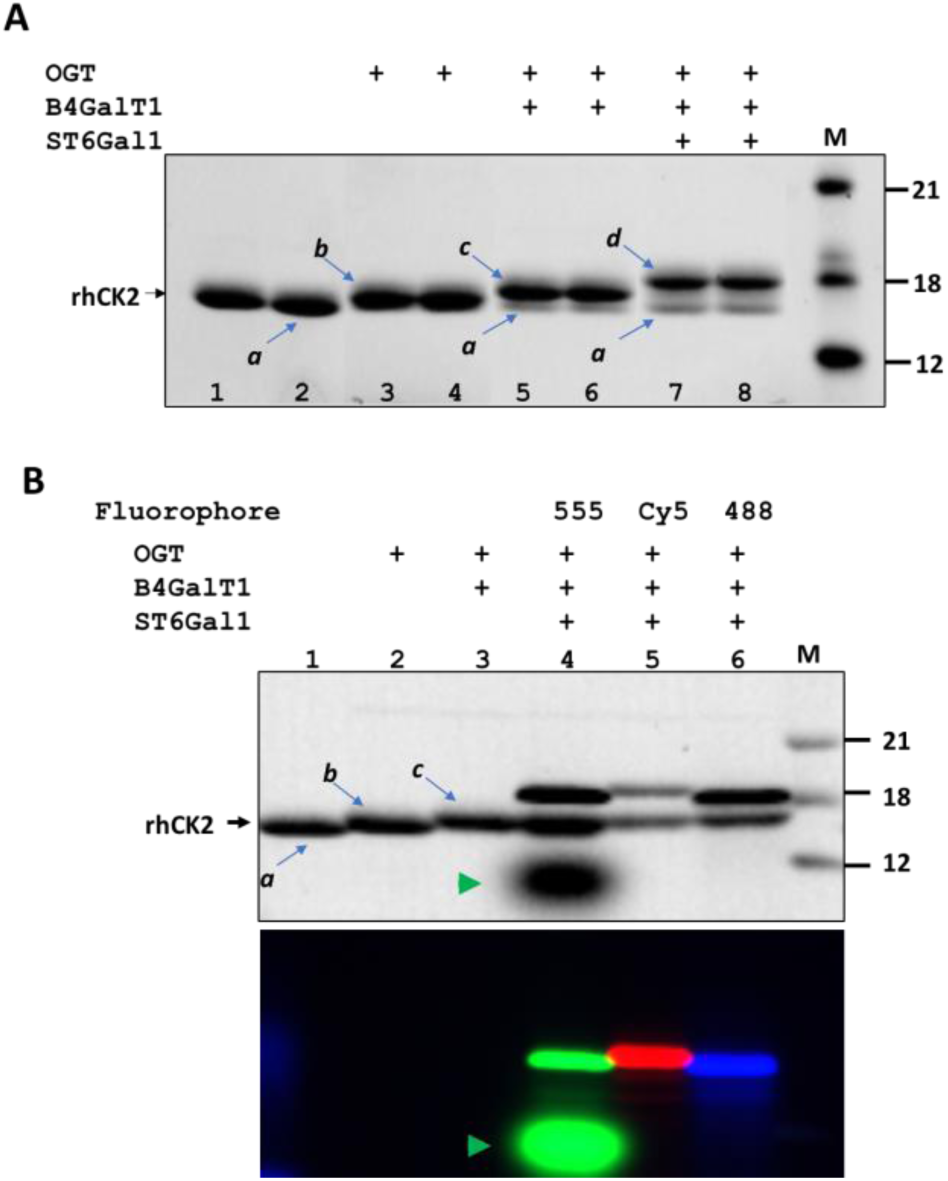
Method validation with recombinant human CK2 (rhCK2). A) *O*-GlcNAc on rhCK2 can be sequentially galactosylated and sialylated. Stepwise modification rhCK2 (labeled as *a*) by OGT, B4GalT1, and ST6Gal1 (their products are labeled as *b, c*, and *d*, respectively) resulted in successive mobility shift on SDS-PAGE that was visualized by trichloroethanal (TCE) imaging. B) Labeling *O*-GlcNAc on rhCK2 with different fluorophore-conjugated sialic acids. *O*-GlcNAc was first introduced to rhCK2 and then labeled by different fluorophore-conjugated sialic acids. Samples were separated on SDS-PAGE and visualized with TCE imaging (upper panel) and fluorescent imaging (lower panel). From lane 4 to 6, *O*-GlcNAcylated rhCK2 was labeled with AlexaFluor 555 (555), Cy5, and AlexaFluor 488 (488), respectively. Free AlexaFluor 555 dye was not removed in lane 4 (marked by a green arrow). M, molecular marker (range in kDa).

The method of fluorescent detection of *O*-GlcNAc with tandem labeling was first tested on the nuclear extract of HEK293 cells (Fig. 3). Labeling of the extract resulted in numerous fluorescent bands across the SDS-gel. Furthermore, these bands were obliviated by OGA pretreatment and augmented by OGT pretreatment, confirming that the signals were due to *O-*GlcNAcylation. The results not only confirm the abundance of *O*-GlcNAcylation on nuclear proteins but also suggest that the labeling method is applicable to detect *O*-GlcNAc on unknown proteins. The method was also validated on various purified proteins (Supplemental Fig. 3). As expected, AKT1, PFKFB3 ^22^, PFKFB4, and PKM ^28^ that are known to be targeted for *O*-GlcNAcylation were modified by OGT and subsequently labeled by Cy5.

**Figure 3.**
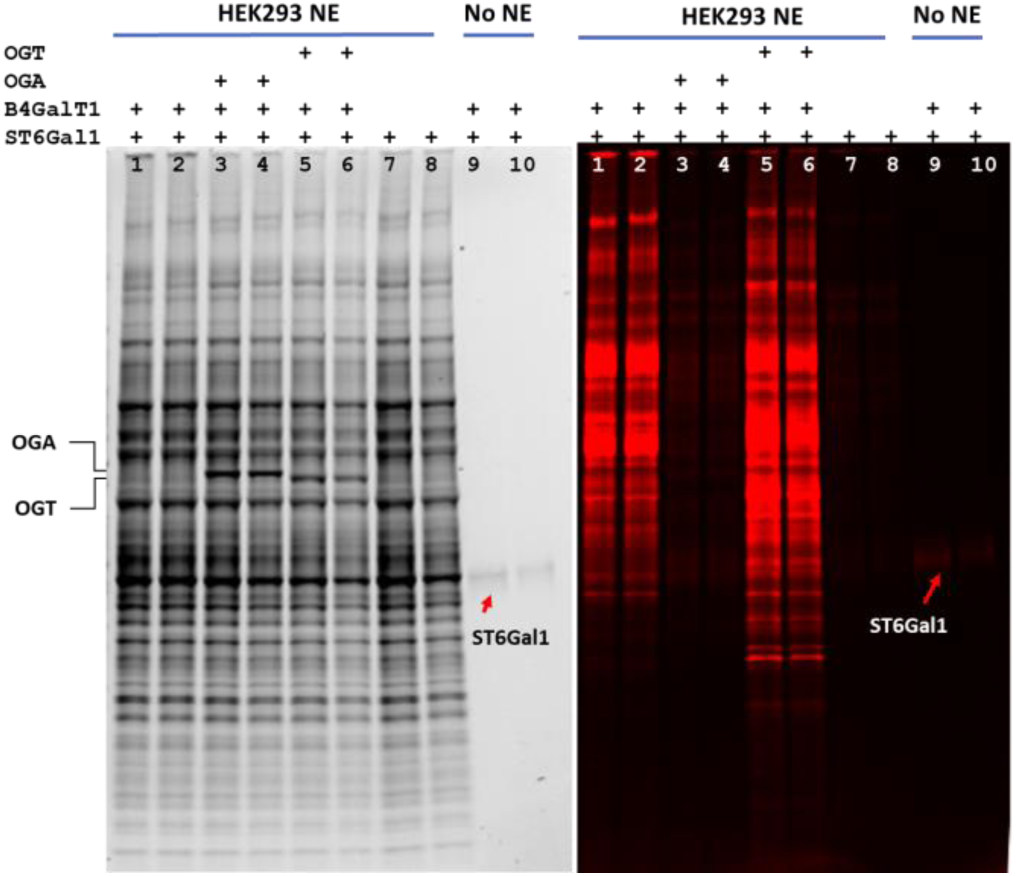
Detection of *O*-GlcNAc on nuclear extract (NE) of HEK293 cells. *O*-GlcNAc was detected without pretreatment (lane 1 and 2), after OGA treatment (lane 3 and 4), after OGT treatment (lane 5 and 6), in the absence of B4GalT1 (lane 7 and 8). No NE was applied in lane 9 and 10. All samples were separated on an SDS-PAGE and visualized with TCE imaging (left panel) and fluorescent imaging (right panel).

We then applied this method to analyze *E. coli* expressed truncated version of HIF1α (Supplemental Fig. 4) along with some other known targets of *O*-GlcNAcylation including SREBP1c and NOD2 (Fig. 4). To our excitement, it was found that recombinant HIF1α was labeled after OGT treatment, suggesting that HIF1α is a target for *O*-GlcNAcylation.

**Figure 4.**
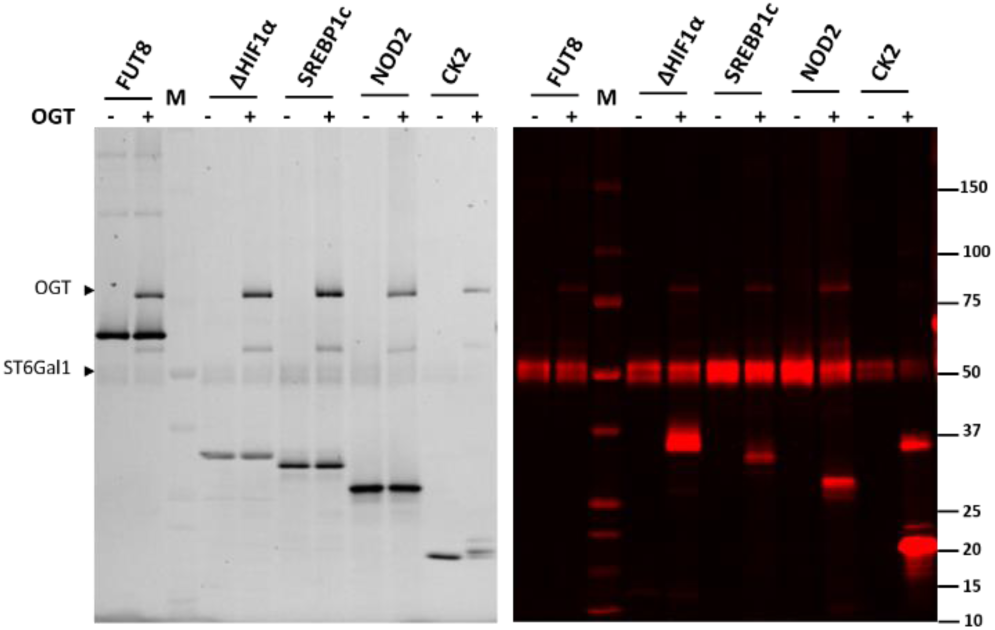
Detection of *O*-GlcNAc on truncated HIF1α (ΔHIF1α). Recombinant SREBP1c, NOD2, and CK2 were probed for *O*-GlcNAc as positive controls with Cy5 before and after OGT treatment and separated on SDS-PAGE and visualized with TCE imaging (left panel) and fluorescent imaging (right panel). FUT8 served as a negative control. OGT and ST6Gal1 exhibited self-modification.

To find out whether HIF1α can be modified by *O*-GlcNAc *in vivo*, full-length HIF1α expressed in HEK293T cells (Supplemental Fig. 4) was purified and probed for *O*-GlcNAc with tandem glycan labeling (Fig. 5). Direct labeling resulted in multiple fluorescent bands above and beneath the position of the unmodified HIF1α but did not when B4GalT1 was absent from the labeling mix or the sample was pretreated with OGA (right panel in Fig. 5), suggesting the presence of *O*-GlcNAc on the sample. Consistent with the positive labeling on the truncated version of HIF1α (Fig. 4 and lane 7 in Fig. 5), OGT pretreatment resulted in significantly increased labeling on a band corresponding to the unmodified HIF1α. Western blotting of the gel with anti-FLAG antibody revealed protein species with a molecular weight higher than that of the unmodified full-length HIF1α (middle panel in Fig. 5), suggesting the presence of ubiquitylated HIF1α. Ubiquitylation is known to lead the mobility shift of HIF1α and its further degradation in SDS-PAGE ^29^, which explains the appearance of the labeled bands above and beneath the unmodified HIF1α. The relative strong labeling on bands above and beneath the unmodified HIF1α suggests that there is a positive correlation between *in vivo O*-GlcNAcylation and ubiquitination on HIF1α.

**Figure 5.**
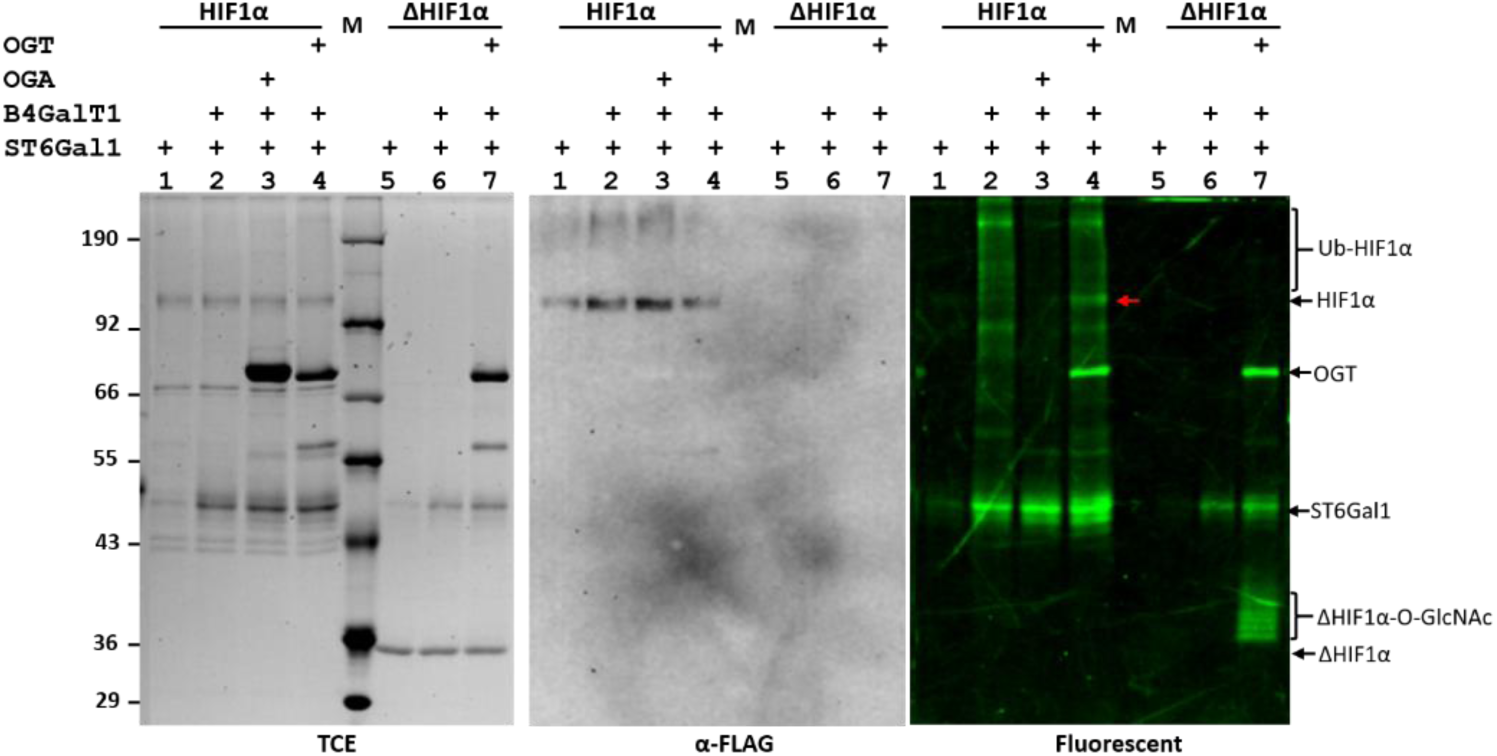
Detecting O-GlcNAc on HEK293T cell-expressed HIF1α. Both HEK293T cell and *E. coli* expressed HIF1α Samples were detected for *O*-GlcNAc with AlexaFluor 555 under indicated conditions. Lane 3 was pretreated with OGA. Lane 4 and 7 were pretreated with OGT. In lane 1 and 5, labeling was performed in the absence of B4GalT1. The gel was visualized by trichloroethanal (TCE) imaging (left panel) and fluorescent imaging (right panel) and then subject to Western blotting with anti-FLAG antibody (middle panel). While OGA pretreatment abolished the labeling, OGT pretreatment increased the labeling on unmodified HIF1α (red arrow). M, molecular marker.

HIF1α is the master regulator in the hypoxia pathway and is activated when the oxygen level is low. An increasing level of HIF1α during hypoxia leads to the upregulation of the glycolysis and hexosamine pathways, therefore, an increased level of *O*-GlcNAcylation, which leads to feedback regulation of the HIF pathway ^15, 30^. However, it is not known whether HIF1α is directly regulated by *O*-GlcNAcylation. Here, we provide evidence that HIF1α can be *O*-GlcNAcylated by OGT *in vitro* and is *O*-GlcNAcylated *in vivo*. HIF1α is best known to be modified by proline hydroxylation ^31, 32^ and subsequent poly-ubiquitination for proteasomal degradation ^29, 33^. The exact biological roles of *O*-GlcNAcylation on HIF1α remains to be investigated. *O*-GlcNAcylation on HIF1α likely counteracts the effect of poly-ubiquitination, therefore positive feedback regulates the HIF pathway, which explains several observations reported previously including that high level of HIF1α correlates to elevated OGT in human breast cancers^15^.

Compared to the previous methods of *O*-GlcNAc detection via click chemistry reaction^22, 34^, the current method has the following advantages or features. First, the method is far more convenient as it eliminates click chemistry reaction and allows direct imaging of samples separated on an SDS gel without the time-consuming membrane transfer and subsequent chemiluminescent detection. Second, the method has eliminated all side effects caused by click chemistry reagents, such as oxidative cleavage of target proteins by copper ions and non-specific labeling observed with copper-free click chemistry reagents. Third, by eliminating the side reactions, the method has also improved the labeling specificity. Although ST6Gal1 can add fluorophore-conjugated sialic acid to other types of glycans, particularly N-glycans^25, 35^, cytosolic and nuclei proteins that are *O*-GlcNAcylated are usually devoid of these glycans. In addition, specificity under questioning can always be confirmed with additional OGA and OGT pretreatment. Fourth, the method is expected to have high sensitivity. Since femtomole level of Cy5-labeled glycans can be measured^26^, it is reasonable to expect that a same level of *O*-GlcNAc can be detected.

## ASSOCIATED CONTENT

### Supporting Information

Experimental procedures, amino acid sequences of recombinant CK2 and HIF1α, mass spectrometry analysis of the B4GalT1, and ST6Gal1 modified rhCK2, fluorescent detection of O-GlcNAc on metabolic enzymes are included in the supporting information.

### Notes

The authors declare no competing financial interests.

## ACKNOWLEDGMENT

We thank Hai-Bin Ruan for critical reading of the manuscript. This work is supported by R&D Systems, a Bio-Techne brand, and by National Institute of Health (R35GM124896 to Y.C).

## Supplemental Information

### Materials and Method Details

#### Materials

Recombinant CK2, B4GalT1, ST6Gal1, PGK1, HK1, HK2, AKT1, FUT8, HIF1α, SREBP1c, NOD2, G6PD, PFKFB3, PFKFB4, PGK1, and PKM were from R&D Systems, Bio-techne. Alkyne-Alexa Fluor® 488, alkyne-Alexa Fluor® 555 were from Thermo Fisher Scientific. Cy5-alkyne, MG132, Flag-Beads and FLAG peptide were purchased from Sigma-Aldrich. Fluorophore-conjugated CMP-Sialic acids were prepared as previously described ^1^.

#### Sample Pretreatment Reactions

For OGT pretreatment, a sample (< 10 μL) mixed with 1 μg of OGT and 10 nmol UDP-GlcNAc was diluted in the glycosyltranserase buffer (25 mM Tris pH 7.5, 10 mM of MnCl_2_ and 10 mM CaCl_2_) to 20 μL and then incubated at 37°C for 30 minutes. For OGA pretreatment, a sample (< 10 μL) mixed with 1 μg of OGA and diluted to 20 μL with 50 mM MES pH 5.5 was incubated at 37°C for 30 minutes.

#### Tandem Fluorescent Glycan Labeling on O-GlcNAc and Imaging

For a typical labeling reaction, the above pretreated sample or 1 to 5 μg of a target protein was mixed with 0.2 μg ST6Gal1, 0.2 nmol fluorophore-conjugated CMP-Sialic acid, 0.2 μg B4GalT1, 10 nmol UDP-Gal and diluted to 40 μL in the glycosyltranserase buffer. The mixture was then incubated at 37°C for 30 minutes. The reaction was then separated by sodium dodecyl sulfate–polyacrylamide gel electrophoresis (SDS-PAGE) and the gel was imaged by both a fluorescent imager FluorChem M (ProteinSimple, Bio-techne) and a UV imager through trichloroethanol (TCE) staining ^2^.

#### *In vivo* purification of full-length HIF1α

HEK293T cells transfected with Flag-HIF1α plasmid (HG11977-NF, Sino Biological) were plated in 10 cm petri dishes for 12 hours and then followed by hypoxia treatment (1% O_2_) for another 12 hours. The cells were treated with 5 μM MG132 (Sigma Aldrich) 4 hours before harvesting. For FLAG immuno-precipitation, cell lysates were prepared with a cell lysis buffer (150 mM NaCl, 50 mM Tris-HCl at pH 7.5, 10% glycerine, 0.5% NP40) supplemented with protease inhibitor (cOmplete™ Protease Inhibitor Cocktail, Roche). Each cell lysate was then incubated with 40 μL of Flag-Beads (A2220, Sigma Aldrich) overnight and washed with the cell lysis buffer four times and then eluted with 90 μL 150 ng/μL 3X FLAG peptide (Apexbio.com).

#### Cell Extract Preparation for HEK293 Cells

HEK293 cells were harvested and gently suspended using 600 μL of an isotonic lysis buffer (10 mM Tris–HCl 7.5, 2 mM MgCl_2_, 3 mM CaCl_2_, 0.3 M sucrose, 0.2 mM PMSF and 0.5 mM DTT) and incubated for 15 min on ice. After centrifugation for 5 min at 420 *g*, the pellet was resuspended with 150 μL of the isotonic lysis buffer. The cells were then slowly passed through a syringe with a 27gauge needle 6 times. The whole lysate was centrifuged for 20 min at 11,000 *g* and the supernatant (cytoplasmic fraction) was collected. The pellet was briefly washed with the isotonic lysis buffer and then extracted using 150 μL extraction buffer (10 mM 4-(2-hydroxyethyl)-1-piperazineethanesulfonic acid, pH 7.9, 1.5 mM MgCl_2_, 0.2 mM ethylenediaminetetraacetic acid, 25% Glycerol, 0.42 M NaCl, 0.2 mM PMSF, and 0.5 mM DTT). The lysate was occasionally vortexed and incubated on ice for 30 min. The supernatant (nuclear fraction) was harvested after centrifugation at 21,000 *g* for 5 min. Extracts were stored at −80°C.

#### Mass Spectrometry Analysis of Recombinant CK2

Recombinant human CK2 was successively treated with OGT, B4GalT1 and ST6Gal1 in the presence of their natural donor substrates and were analyzed with mass spectrometry. Electrospray ionization mass spectrometry (ESI-MS) analysis on the glycosylated CK2 peptide was performed using a Thermo Scientific Q Exactive HF Mass Spectromerter coupled to a Thermo Vanquish system. Samples (10 μL) were separated on a HiChrom Vydac 214MS C4 column (50 x 2.1 mm, 5 μm) with a Vydac 214MS C4 guard column using a gradient flow set at 400 μL/min. Mobile phases consisted of A – 0.05% trifluoroacetic acid in water and B – 0.05% trifluoroacetic acid in acetonitrile. The gradient was 0-2 minute – 98% A, 2-7 minutes – 98-2% A, 7-7.5 minutes – 2% A, 7.5-8 minutes – 2-98% A. Total run time was 12 minutes. The mass analyzer was run in full scan from m/z 600 - 3000 operating in positive mode. Full MS parameters: no in-source CID; 3 microscans; resolution = 240,000; AGC target = 5e6; Maximum IT = 200 ms; Number of scans = 1, Profile spectrum. Data was deconvoluted using Thermo Bio-Pharma Finder software.

---

**Supplemental Figure 1.**
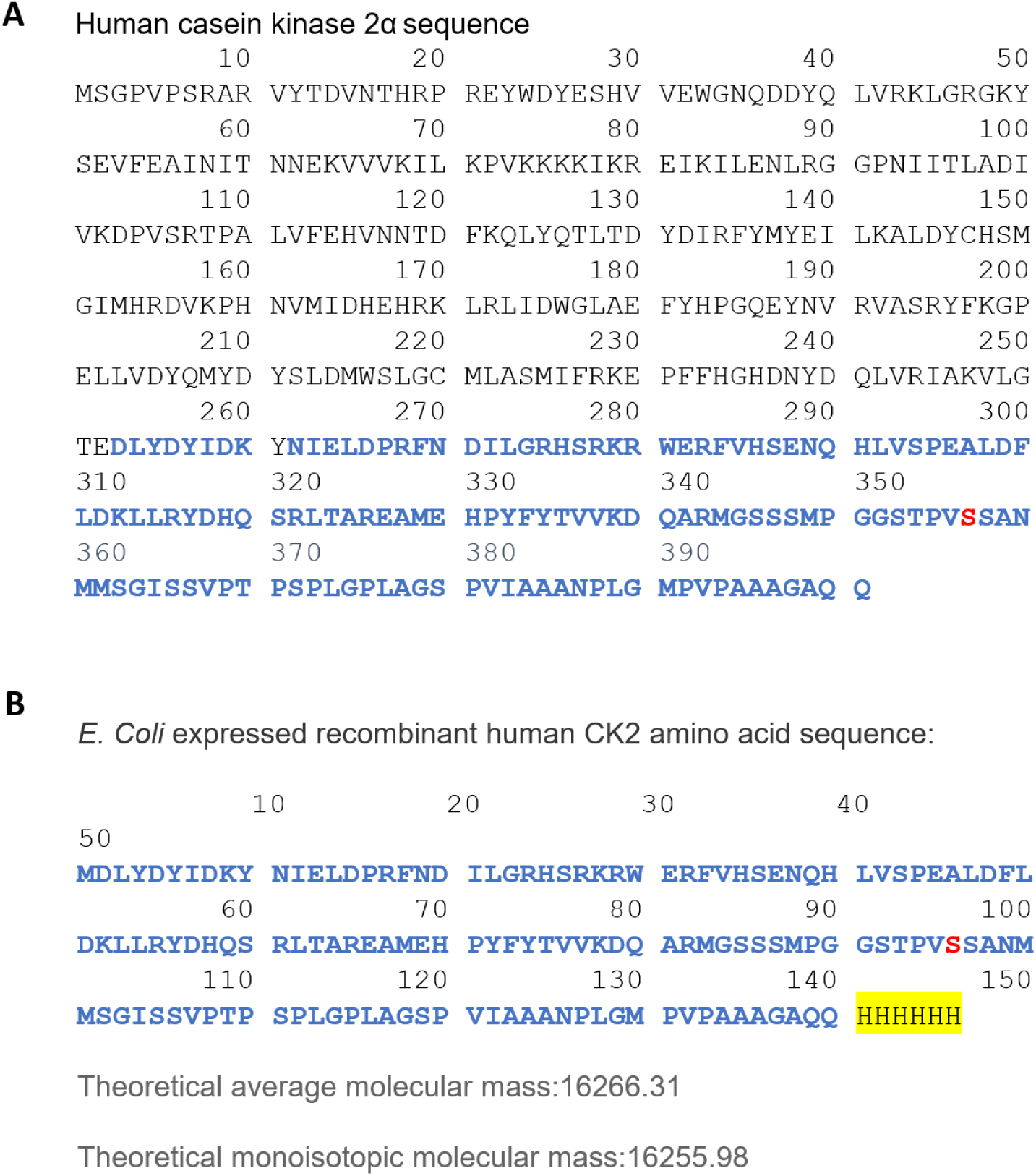
Sequences of human and E. Coli-expressed recombinant CK2. A) Sequence of human casein kinase 2α (CK2) (Uniprot P68400). B) The sequence of *E.coli* expressed recombinant CK2 (rhCK2, R&D Systems Cat# 7957-CK). A 6x His tag was added to the C-terminal of the recombinant CK2. The O-GlcNAc site at Ser347 is highlighted in red.

**Supplemental Figure 2.**
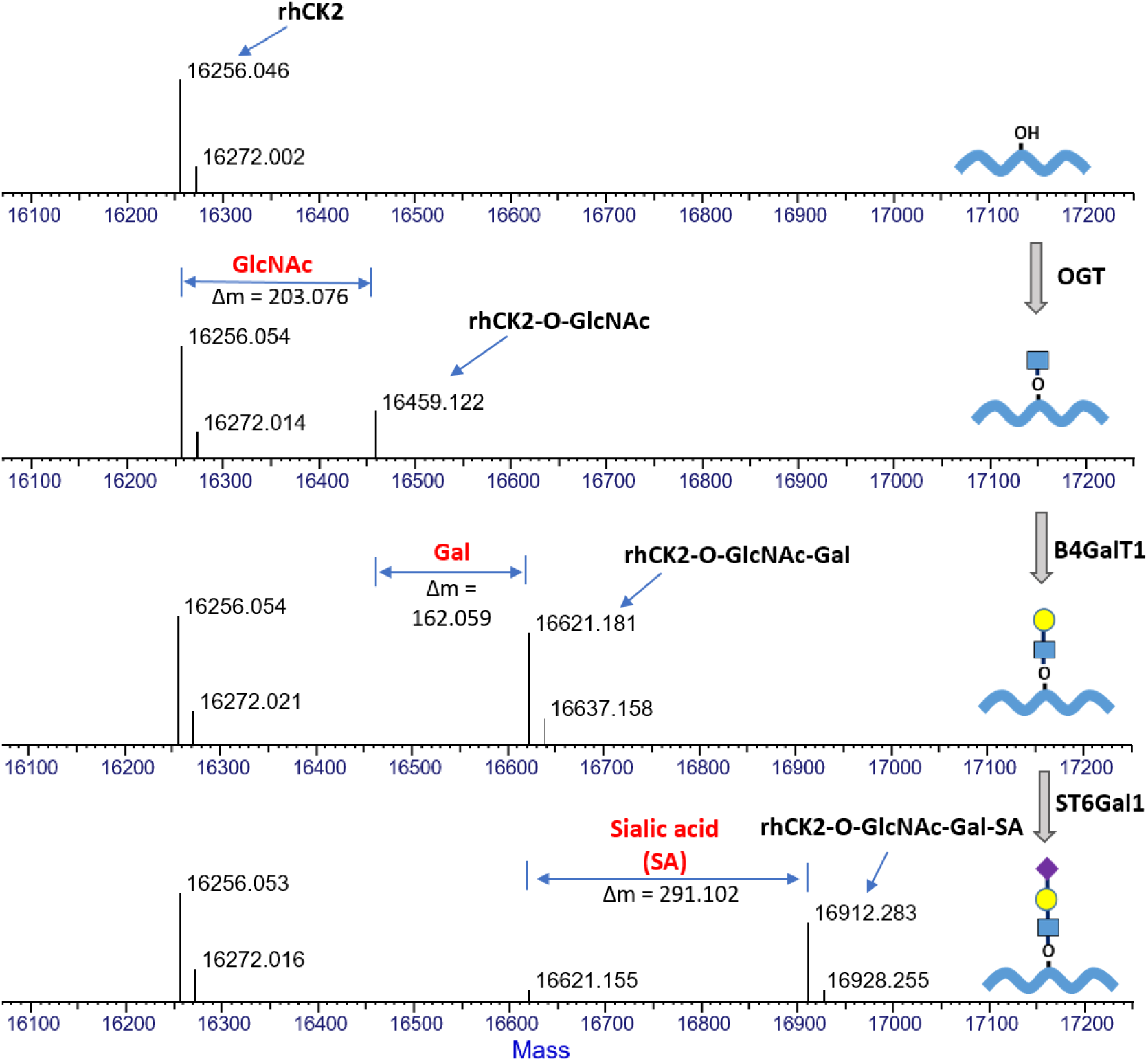
*O*-GlcNAc on recombinant CK2 can be sequentially galactosylated and sialylated. Stepwise modification of recombinant CK2 by OGT, B4GalT1, and ST6Gal1. The products were analyzed by mass spectrometry analysis.

**Supplemental Figure 3.**
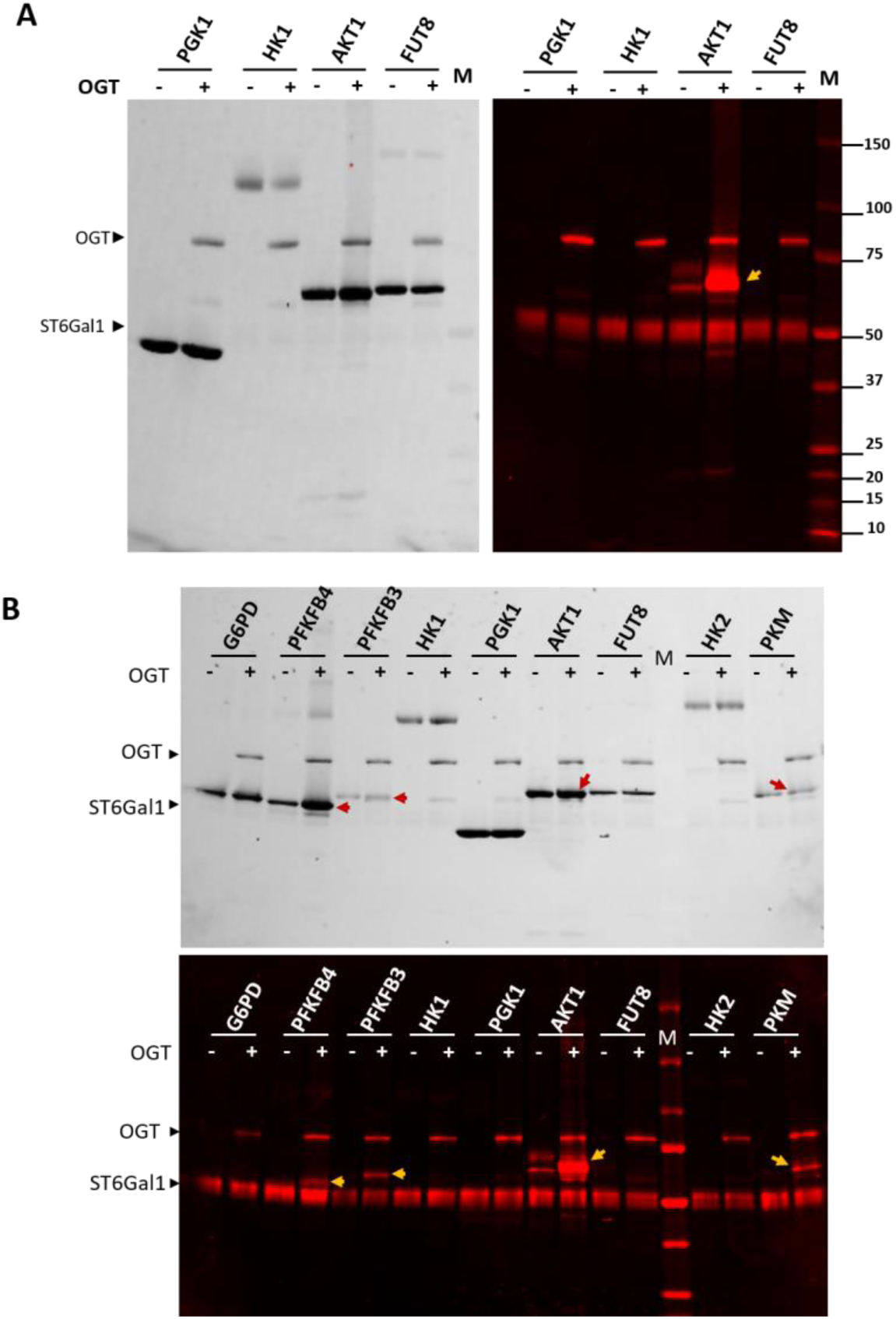
Detection of *O*-GlcNAc on various recombinant enzymes. A) Group I enzymes and B) Group II enzymes were probed for *O*-GlcNAc before and after OGT treatment. The indicated enzymes (around 1 μg per lane) were labeled with Cy5-conjugated sialic acid. FUT8 is a Golgi resident enzyme and was used as a negative control. Labeling enzyme ST6Gal1 showed significant self-labeling. AKT1 expressed in sf9 baculovirus exhibits lower level of O-GlcNAcylation without OGT treatment. *In vitro* O-GlcNAcylation (indicated with yellow arrows) was observed on PFKFB3, AKT1, and PKM. G6PD, Glucose-6-phosphate dehydrogenase. PFKFB3, 6-phosphofructo-2-kinase/fructose-2,6-biphosphatase 3. PFKFB4, 6-phosphofructo-2-kinase/fructose-2,6-biphosphatase 4. HK1, Hexokinase-1. HK2, Hexokinase-2. PGK1, Phosphoglycerate kinase 1. AKT1, RAC-alpha serine/threonine-protein kinase or protein kinase B (PKB). FUT8, α1,6-fucosyltransferase. PKM, pyruvate kinase muscle isozyme. All labeled samples were separated on SDS-PAGE and visualized with TCE imaging and fluorescent imaging. OGT and ST6Gal1 exhibited self-modification and were indicated.

**Supplemental Figure 4.**
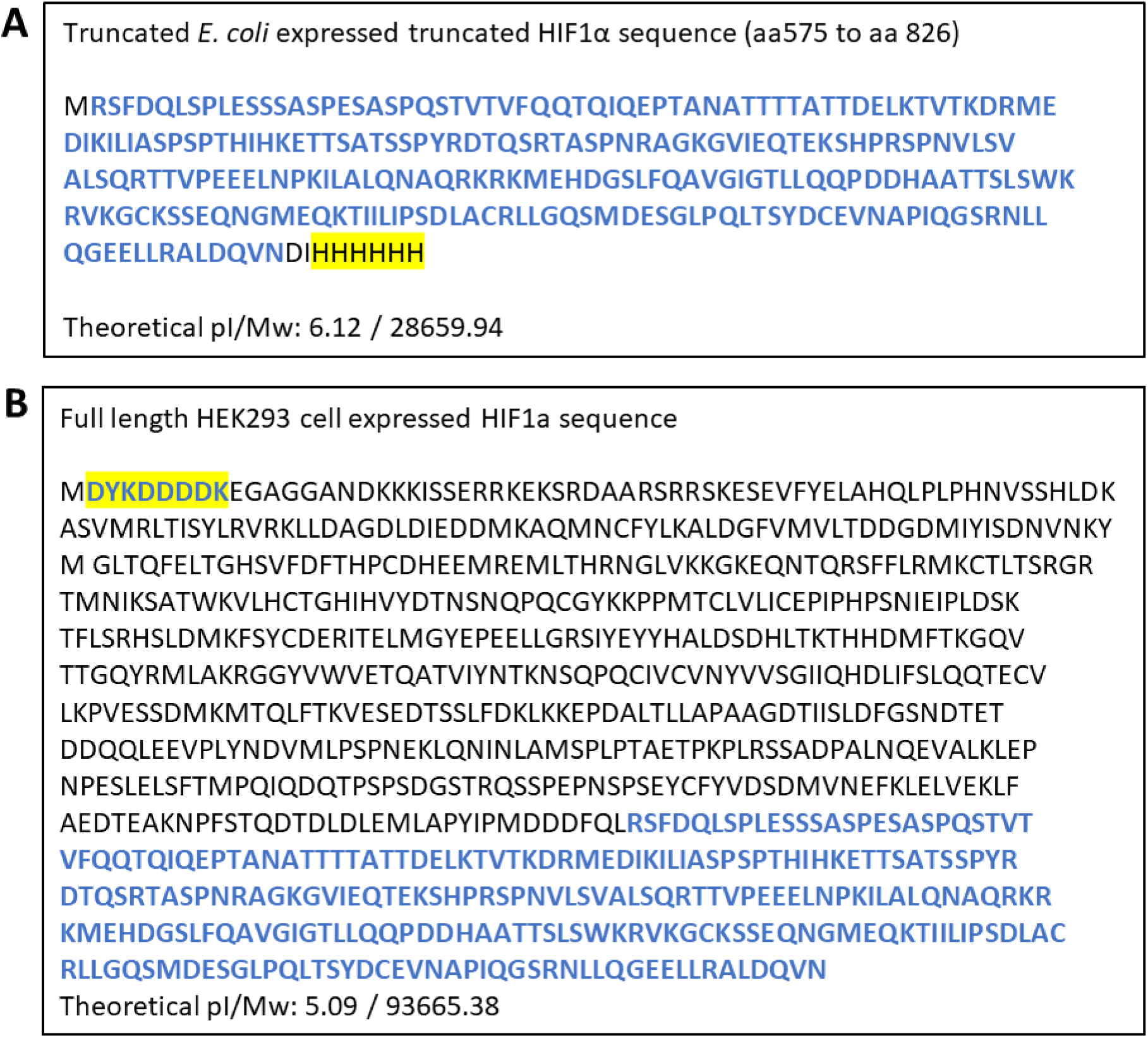
Sequences of recombinant and FLAG-tagged human HIF1a. A) The sequence of the truncated version of HIF1α with a 6xHis-tag expressed in *E. coli.* B) The sequence of the full-length version of HIF1α with a FLAG-tag expressed in HEK293 cell. The sequence that is common to both expressed versions is highlighted in blue. The 6xHis tag and the FLAG-tag are highlighted in yellow.

